# Multi-Contrast MRI Inputs Enable Self-Consistent Tissue Segmentation & Robust Perivascular Space Identification

**DOI:** 10.64898/2026.04.03.716409

**Authors:** Jeffrey L. Gunter, Gregory M. Preboske, Benjamin D. Persons, Scott A. Przybelski, Christopher G. Schwarz, Audrey Low, Prashanthi Vemuri, Ronald C. Petersen, Clifford R. Jack

**Author notes:** Correspondence: Jeffrey L. Gunter, Ph.D., Department of Radiology, Mayo Clinic, Rochester, MN, USA.

## Abstract

Different MRI image contrasts are designed to highlight various tissue properties and combining them allows extension of probabilistic segmentation beyond the commonly used “gray-white-CSF” models. This work describes a fully automated method that combines T1-weighted, T2-FLAIR, and conventional T2-weighted images to provide internal consistency across prediction of tissue segmentations including segmentation of superficial and deep gray matter, white matter hyperintensities, and MR-visible perivascular spaces. Results from 773 imaging datasets from 403 participants in the Mayo Clinic Study of Aging and Mayo Clinic Alzheimer’s Disease Research Center (ADRC) are presented.

## Introduction

Automated tissue segmentation (classification) of structural MR images has a long history (e.g. Wells et al., 1996; Held et al., 1997; reviewed in Ashburner (2012)). Segmentation of white matter hyperintensity (WMH) on standard T2-weighted and T2-FLAIR images also has a long history, including early works such as Jack et al. (2001). Many deep learning methods have been developed for both WMH identification and tissue classification (e.g., Ghafoorian et al., 2017; Li et al., 2018; Ronneberger et al., 2015; Wachinger et al., 2018; Rahmani et al., 2024). Identification of MR-visible perivascular spaces (PVS), generally on T1-weighted or T2-weighted imaging, is of increasing interest as it may be a marker of glymphatic system function and/or a marker of cerebrovascular risk (Wardlaw et al., 2013; Iliff et al., 2012; Potter et al., 2015). Although all these methods address the same gross anatomy, the methods are not generally combined in a self-consistent fashion. To address this, we leveraged the classic SPM Unified Segmentation (Ashburner & Friston, 2005) approach, extending its rarely used ability to segment with multiple input contrasts to combine them in a self-consistent, fully automated framework.

Segmentation identifying gray matter (GM), white matter (WM), and cerebrospinal fluid (CSF) using T1-weighted images is common. Multiple frameworks are readily available with SPM12 unified segmentation, FreeSurfer, and FAST as prime examples (Ashburner & Friston, 2005; Fischl, 2012; Zhang et al., 2001). Methods supporting automated delineation of WMH are also available, largely using T2-FLAIR images as input (DeCarli et al., 2005; Schmidt et al., 2012). Machine Learning / Deep Learning methods for WMH segmentation are an area of active development (reviewed in Rahmani, 2024). Methods supporting automated delineation of PVS using T1-weighted and/or T2-weighted or proton density weighted images exist (e.g., Ballerini et al., 2016; Boespflug et al., 2018; Ballerini et al., 2018; Schwartz et al.,2019; Sepehrband et al., 2019; Rashid et al., 2023; Zhang et al., 2024). The use of multiple MRI contrasts within a unified probabilistic framework has been well established in the context of multiple sclerosis lesion segmentation, where combining T1-weighted and T2-FLAIR images improves discrimination of lesion tissue from normal-appearing white matter (Shiee et al., 2010; Schmidt et al., 2012).

Combining T1-weighted images, T2-FLAIR, and T2-weighted images provides a broader range of input information than any single image alone. For example, T1-weighted images have superior contrast between GM and normal appearing WM compared to typical T2-FLAIR images, while the fluid attenuation in T2-FLAIR strongly differentiates CSF and diseased white matter. Conventional T2-weighted images provide high contrast between GM and CSF as well as CSF and dura. SPM Unified Segmentation has always supported multiple contrasts as input, though this functionality appears seldom used. The additional information from combining multiple input contrasts allows us to expand the space of output tissue classes beyond GM, WM and CSF. Specifically, we have superficial GM, normal appearing white matter (NAWM), CSF, deep gray matter (DGM) and WMH classes. The resulting contrast model parameters can be exploited in a subsequent step to isolate PVS.

## Methods

### Overview

Structural MRI contrasts are chosen to highlight different aspects of anatomy. By combining T1-weighted, T2-weighted and T2-FLAIR images, multiple tissue classes may be better understood compared to classification from any sequence in isolation. The model, as previously mentioned, identifies superficial gray matter (GM), normal appearing white matter (NAWM), CSF, deep gray matter (DGM), WMH, and PVS classes. Nuisance classes for skull, scalp and air are also included. Table 1 outlines the qualitative appearance of each tissue class on the various images.

**Table 1:**
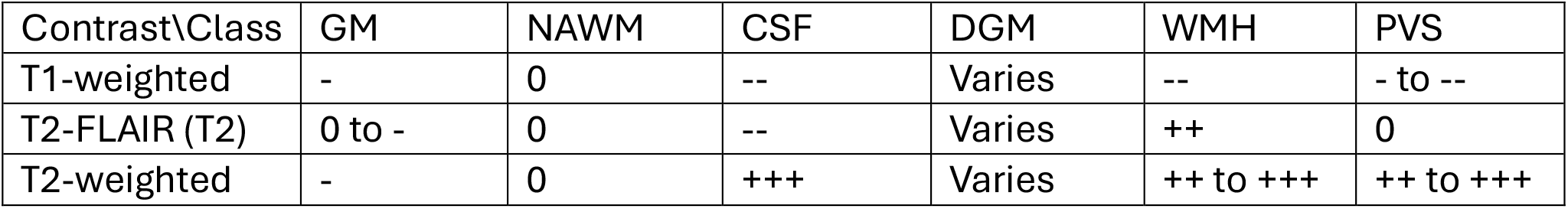
Qualitative description of relative contrasts for tissue classes in different MRI input images. Normal appearing white matter (NAWM) is chosen as the reference (0) for each contrast (row) with +/-indicating higher and lower relative intensities. Deep gray matter is highly variable between individuals.

### Imaging Data and Participant Selection

Imaging data from 773 MR studies of 403 study participants were used to demonstrate the feasibility and performance of extended segmentation and PVS identification. Participants were drawn from the Mayo Clinic Study of Aging (MCSA, a population based study of cognitive aging) and the Mayo Clinic Alzheimer’s Disease Research Center (ADRC, a behavioral neurology clinic -based cohort). All imaging was done on a Siemens 3T Prisma scanner using a 64 channel head coil. The imaging protocols are similar to the protocol used in ADNI-3 and ADNI-4. That is, images are 3D acquisitions with roughly isotropic voxel sizes in the range of 1mm x 1mm x 1mm. Details are provided in supplemental materials. We included 325 MCSA study participants who were randomly selected from the larger MCSA cohort to fill 5-year age bins from age 30 to 90 years and a single bin for ages over 90 years with equal numbers of females and males. We also included 78 demented ADRC participants to ensure our sample would have a wide dynamic range in volumetric measurements. A subset of 249 of these participants had longitudinal imaging. A subset of 125 MCSA participants were assessed for PVS. Participants are further described in Table 2.

**Table 2.**
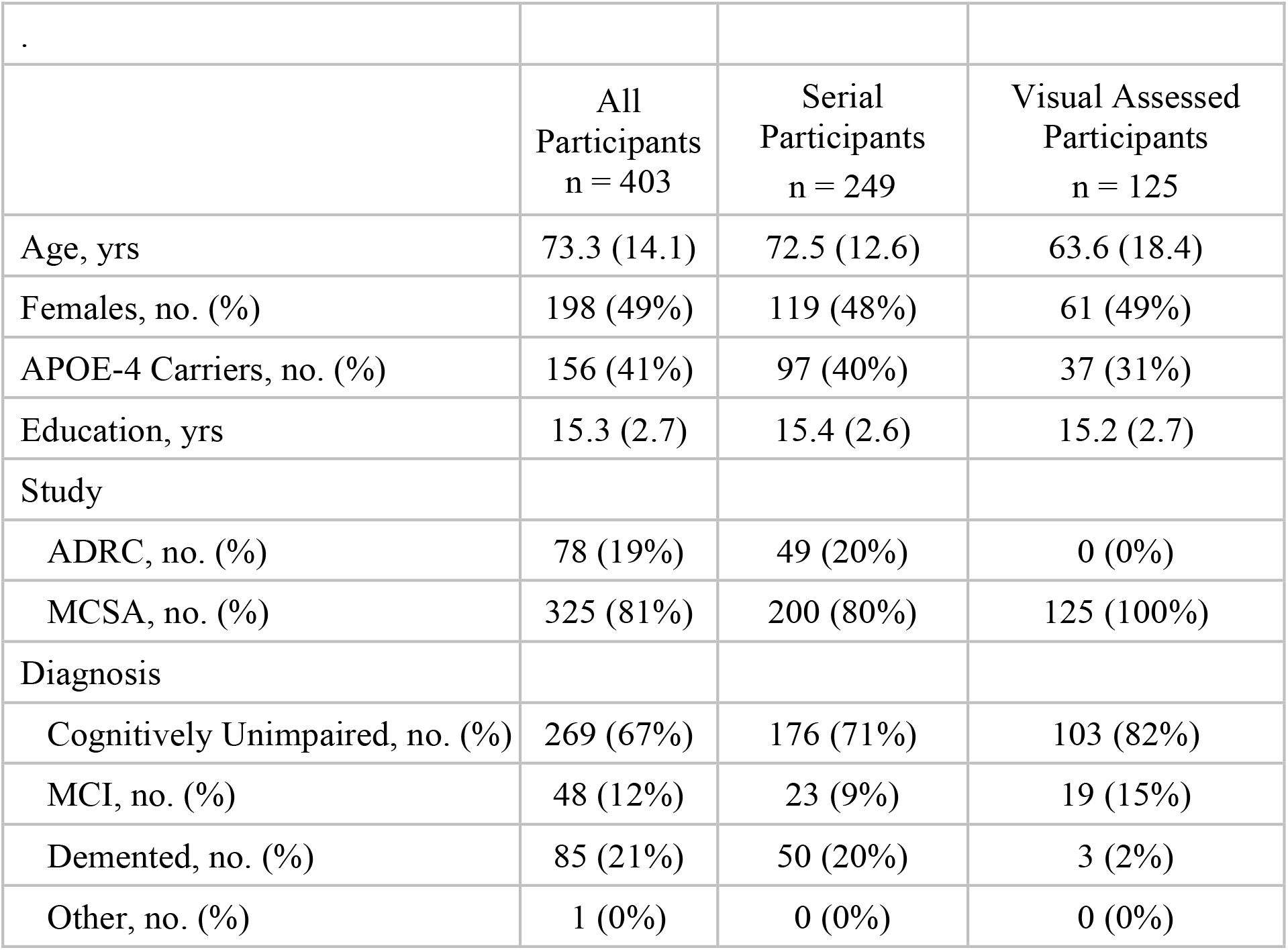
Baseline characteristics table of groups with the mean (SD) listed for the continuous variables and count (%) for the categorical variables.

### Image Processing

The image processing method implemented includes three main components: pre-segmentation image registration, segmentation model estimation, and post-estimation tissue classification and PVS estimation. We used the Mayo Clinic Adult Lifespan Template (MCALT, https://www.nitrc.org/projects/mcalt/) family of images, atlas parcellations and utility masks, and the MCALT single-contrast segmentation model (Schwarz et al. 2016) was the starting point that we extended for multi-contrast. We extended the tissue probability maps (priors) as necessary to include DGM and WMH classes.

### Pre-segmentation Steps

Processing begins with an automated check to ensure that each acquired image set is 3D and has sufficient coverage to capture an adult head, which we assume to be at least 160mm in any dimension. Image coordinate system origins are automatically sanity checked. Any coordinate system origin outside the field of view is set to the center of the field of view. Registration is performed using a swarm of 5 different starting transformations with the spm_coreg function and also one call to reg_aladin from the NiftyReg (Modat et al., 2014) package to determine the best affine transformation between the T1-weighted image to be segmented (subject space) and the T1-weighted average grayscale MCALT image. Spm_coreg returns a log-likelihood value after minimization; the log-likelihood value for the transformation returned by reg_aladin is calculated using the internal function fo spm_coreg. The transformation with best log-likelihood value is chosen. This registration scheme was developed over many years and is extremely robust (i.e. failure rates less than 1 per 10,000 uses). A mask containing the entire head and dilated approximately 10mm is transformed from template into subject space. The T2-weighted and T2-FLAIR images are then registered to subject T1-weighted image space allowing rotation, translation and scaling (9 DOF) with the mask used to focus on the head and ignore differences in neck position and whether shoulders happen to be included in the fields of view. Grayscale T2-weighted and T2-FLAIR images are resampled into the raster of the T1-weighted image using cubic spline interpolation. We thus have three images in the T1-weighted image space and a transformation matrix mapping that space to the MCALT T1-weighted average.

### SPM12 Segmentation Model Estimation

SPM12 unified segmentation determines the deformation between tissue prior probability maps in template space and subject space, bias fields for each input contrast, and a model of tissue contrasts. Empirically, estimating the full desired contrast model in addition to deformation and bias fields de novo is unreliable. This is largely due to anatomic variability. An iterative procedure is adopted: initial segmentation is performed using only the default GM, WM, CSF, and nuisance classes; a DGM class is then added and model parameters are updated; finally, WMH is added and model parameters are again updated. As a 9DOF registration to template space has already been performed, it is provided to the initial segmentation as a starting transformation. Minor modifications to the SPM12 unified segmentation code available with the MCALT package are necessary to enable updating existing model parameters instead of starting from scratch. As contrast classes are added the number of classes in the tissue prior probability maps is also increased. The expanded DGM prior was created by smoothing the relevant regions in the MCALT ADIR122 Atlas. The WMH prior was determined from the smoothed voxel-wise average of WMH maps from a prior unpublished MCSA analysis using a semi-manual method and adding a small flood-fill probability over all brain tissue. The WMH prior loads heavily around the lateral ventricles.

A more detailed description including practical aspects used in the iterative segmentation scheme follows:

- Unified segmentation supports the inclusion of a mask restricting voxels used in model estimation. A mask, propagated from template space that includes the total intracranial volume (TIV) dilated by 20mm to include skull, scalp and air is provided as an input. Analogous to the use of a mask during registration this avoids including shoulders and areas with known severe shading artifacts.
- After each segmentation optimization pass, contrast model parameters, bias-correction fields, and the deformation mapping template and subject coordinates are available. Extending the model includes the following steps. The current estimated bias fields are applied to create temporary bias-corrected images. The current deformation is applied to deform a set of tissue priors including the new class being added. Voxels in regions of high priority for the class being added are identified. Percentile values of the bias-corrected image intensities in those voxels are used to seed an expanded segmentation model intensity parameter set. We are using the default mixture of Gaussians contrast model. That is, we set the mixture of Gaussian centroids using the previously estimated bias-field and deformation. The covariance matrix is seeded with variance values for each tissue class. A subsequent pass through the unified segmentation code with the expanded parameter model and expanded set of prior probability maps is then made to refine all the parameters.

### Post-segmentation

After the segmentation model estimation is complete, several steps are performed to generate directly useable outputs including PVS delineation.

#### Discrete Class Map (argmax) Production

Class probability maps are generated from the final segmentation model. The default SPM12 Markov Random Field smoothing is used. The WMH probability map is refined by requiring the T2-FLAIR intensity be more than two standard deviations above NAWM mean, with the rejected WMH probability being moved into the NAWM class. The argmax operator is applied to the probability maps to assign each voxel to the class with highest probability. We refer to this as the “argmax” segmentation image.

#### PVS identification

PVS identification leverages the segmentation model. Bias-corrected images are created. A rough mask of the supratentorial tissue is warped from the MCALT space into subject space. Voxels with high segmentation output GM probability or near the surface of the lateral ventricles are excluded, leaving WM and DGM tissue as a PVS search space. Small holes in the search space mask are filled. A Frangi filter (Frangi et al., 1998; Descoteaux et al., 2008) bank (sigma from 0.5 to 1.5mm in 0.25mm increments, alpha=0.9 and beta=0.9) is passed over the T1- and T2-weighted images to highlight small vessel-like structures. The filter outputs are summed, clipped to a maximum value of one, and smoothed with a 3mm Gaussian kernel. The full set of tissue priors are warped into subject space and resampled using linear interpolation. The CSF prior map is replaced with the smoothed Frangi filter output and priors are normalized so each voxel sums to unity. That is, we have subject-specific priors explicitly existing in the space of the T1-weighted image. Using the likelihood estimation function internal to unified segmentation, the likelihood values are calculated for voxels within the search region using the subject-specific priors. As in the standard approach to tissue classification, the ratio of likelihoods to the sum of likelihoods in each voxel is formed to estimate class probabilities. This allows voxels with CSF-like intensity profiles in areas with high Frangi filter response to be identified. PVS candidate voxels in the frontal, parietal, occipital and temporal lobes are identified as those with the maximum probability among WM, DGM and the new PVS class. The diversity of intensities in the basal ganglia necessitates further refinement. For each of the putamen, globus pallidus, thalamus, and caudate, PVS candidate voxels with intensities below 2.96 times the median absolute deviation (MAD) of T2-weighted image intensities in the respective region are rejected. “MAD times 2.96” is used as a non-parametric measure that maps to approximately two standard-deviations for a normal distribution but is less sensitive to outliers and was chosen empirically. Surviving voxels are clustered using a 26-connected structuring element. Volume and total length of segments allowing for branches are calculated for each cluster. Clusters are also associated to a lobar atlas region based on maximum overlap to avoid splitting clusters across regions. In the argmax tissue classification are voxels now identified with PVS that may have been assigned to a nuisance class or GM during the previous segmentation. These are re-assigned to either NAWM or DGM based on the preponderance of voxels under and immediately adjacent to the PVS as a clean-up step.

As PVS are often assessed in the basal ganglia (BG) and the combined centrum semi-ovale and corona radiata (CSOCR) values from lobar atlas regions may be summed. Note that PVS are common in the external capsule. Voxels in the external capsule may be parcellated into either the basal ganglia or the insula depending on the exact deformation of the lobar atlas regions from template space. To reduce variability related to that we aggregate the insula region into a basal ganglia + insula region.

#### Additional Steps

Given the output of segmentation, TIV, brain, and ventricle masks are constructed using simple morphometry based on the argmax class image. A problem we have commonly encountered in SPM unified segmentation with only a T1-weighted image as input is that the CSF-dura boundary tends to be shifted toward the cortical surface in paticipants with highly atrophic brains. Inclusion of the T2-weighted image provides a high contrast boundary resulting in visually improved TIV masks. Atlases may be warped from template space as necessary using the deformation derived during the SPM12 segmentation step.

### Quality control

First level quality control involves ensuring that the contrast model parameters from segmentation are sensible. As none of the input images are properly quantitative, intensity ratios are used. For example, the CSF:WM intensity ratio for T2-weighted images should be large and less than unity for T1-weighted and T2-FLAIR images. The chief utility of these checks is in the context of processing large collections of data where images may be grossly mis-classified (e.g. a T2-FLAIR identified as a T2-weighted image or a T1-FLAIR image misclassified as a T2-FLAIR). Ideally this should not happen, but it is a simple check when approaching datasets with very large numbers of scanning sessions. The contrast model check found one algorithm failure where the deformation was incorrectly determined despite sensible inputs.

The segmentation model output file includes the filenames for the T1-weighted, T2-weighted, and T2-FLAIR images. The implementation creates files with consistent naming patterns keyed on the T1-weighted input image name. A custom plugin for FSLeyes (McCarthy, 2019) reads the segmentation model (*_seg8.mat) and loads necessary images for review. Specifically, from “bottom to top” the stack of images includes bias-corrected T1-weighted, T2-FLAIR and T2-weighted images, the discrete argmax class identification (color mapped), a PVS map, a representative atlas warped from template space, and the TIV mask. A sample image set is shown in Figure 1.

**Figure 1.**
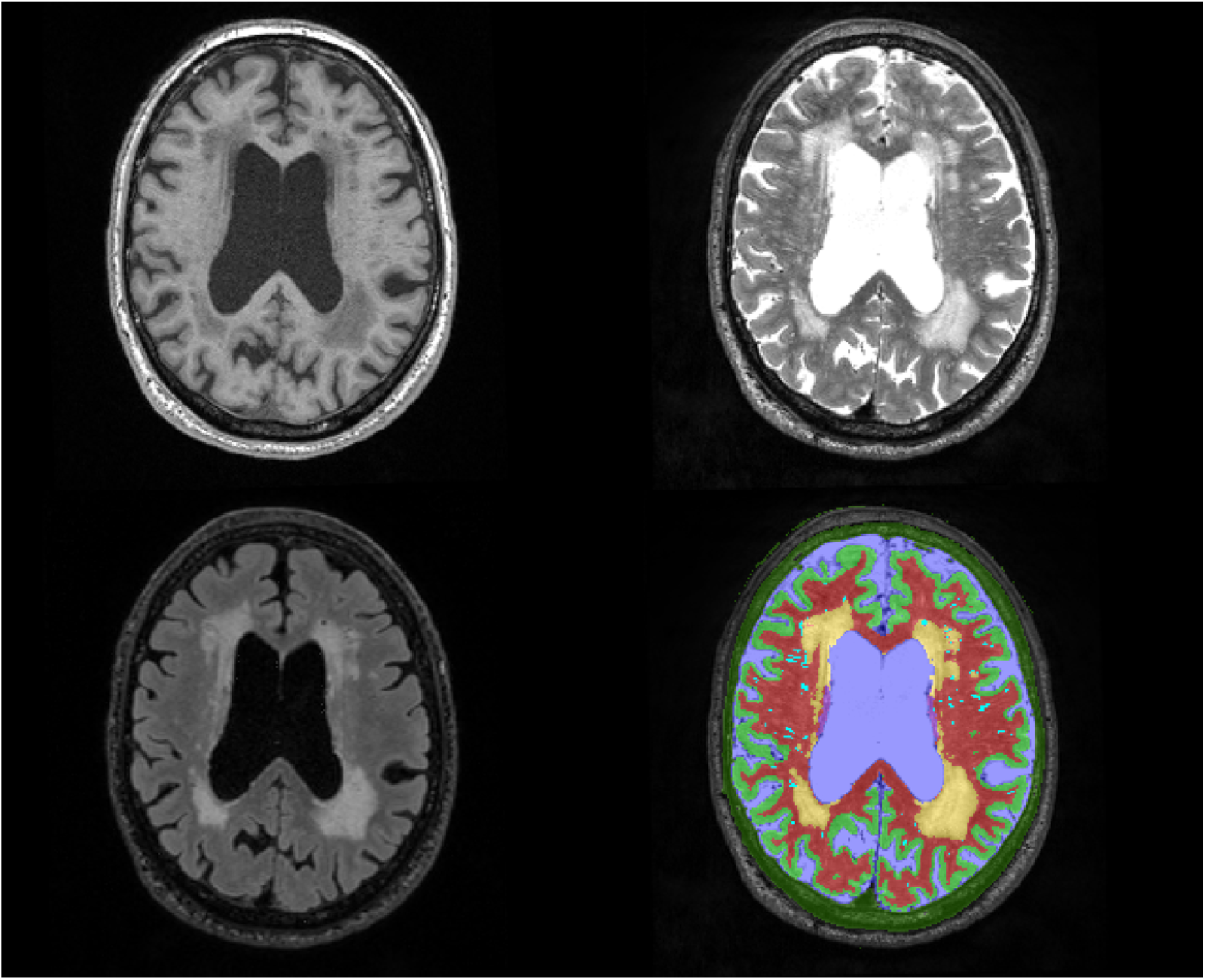
Images from a representative slice from T1-weighted (upper left), T2-weighted (upper right) and T2-FLAIR weighted (lower left) are shown. In the the lower right, the segmentation output is shown overlaid on the T2-weighted image; with CSF in blue, GM in green, NAWM in red, WMH in gold, and the dura nuisance class in a dark green. PVS are shown in a bright blue opaque overlay.

**Figure 1.**
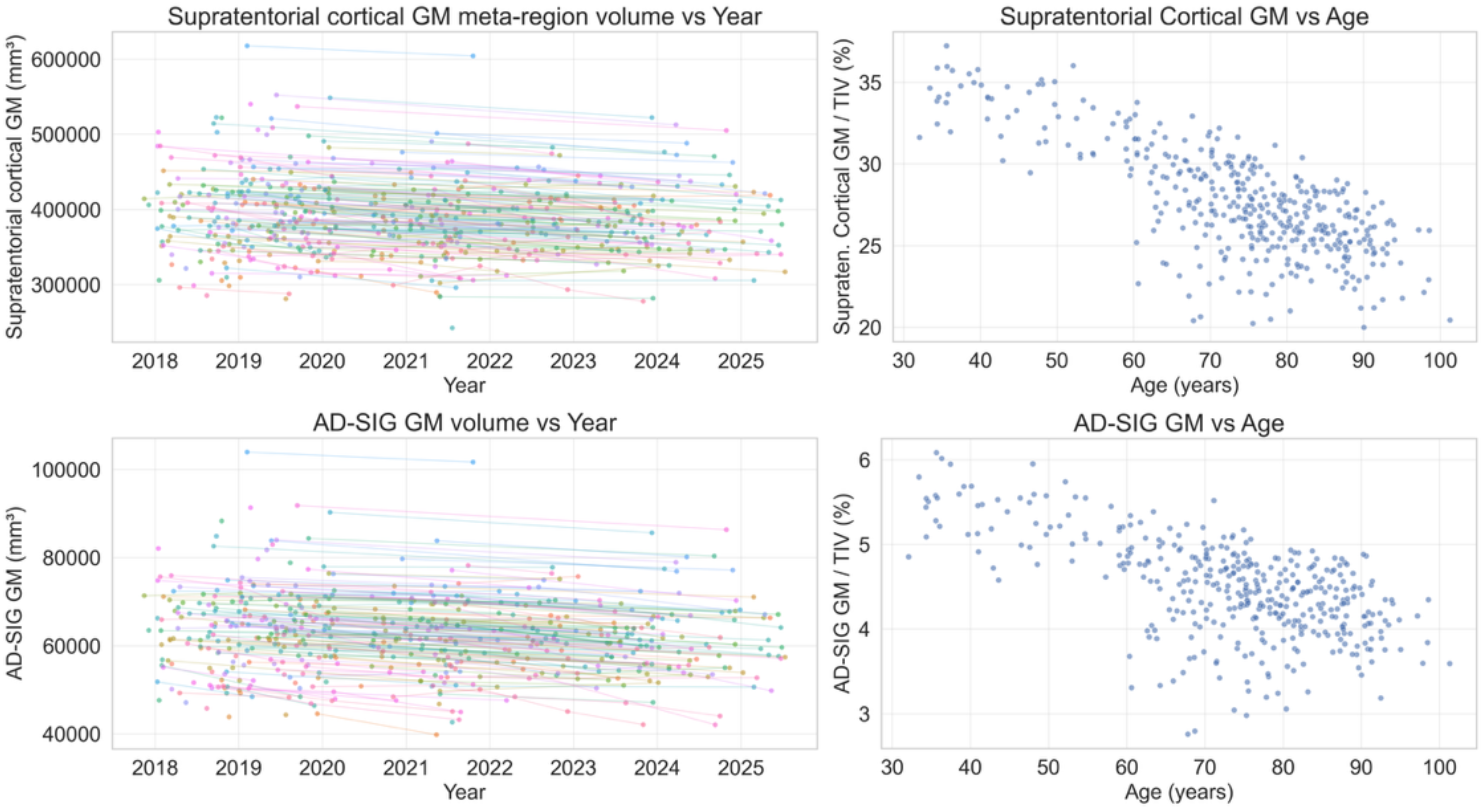
Supratentorial and AD signature (AD-SIG) GM volumes. GM volumes are stable to decreasing over time in individuals with serial exams. GM volume, normalized to TIV, tends to decrease with age. Typical cortical GM in the range of 30% TIV is consistent with, for example, Buckner et al., 2004. The AD-SIG meta-region includes the entorhinal cortex, middle temporal gyrus, inferior temporal gyrus and fusiform gyrus, as these are from a meta region in which cortical thickness is commonly reported Jack et al., 2015.

Overall segmentation quality is checked, with a focus on the GM and NAWM parcellation as pass/fail. WMH and PVS classes assessed on a 5 point scale from underestimated to overestimated, unable-to-assess due to poor image quality or (algorithm) failure. WMH and PVS assessments are graded in the basal ganglia + insula and the CSOCR independently. Details on the QC visualization approach may be found in the supplementary materials.

### Visual PVS assessment

A subset (including n=125 imaging studies) was visually assessed for PVS by two independent raters. The method of Potter et al. (2015) was applied. The purpose of the visual assessment is to provide a comparison with automated results. Visual assessment was performed on a single axial slice in the basal ganglia a single slice in the CSOCR. Slices were selected as the ones with maximum PVS load. On each those slices the hemisphere with high load was evaluated. The numbers of PVS identified visually and the number of automated clusters on those slices and in those hemispheres were counted. Details of visual assessment implementation are provided in the supplemental materials.

#### Validation

Our primary source of external validation for accuracy of segmentation was observations over time within individuals and with age across the whole sample. We assumed the following underlying biological “truths”. TIV would be constant over time within individuals and with age. Measures of brain volumes and brain pathology (WMH and dilated PVS) would decrease/increase respectively over time within individuals and with increasing age.

### Software availability

SPM12 is available at for download at https://www.fil.ion.ucl.ac.uk/spm/software/spm12/. NiftyReg is available on GitHub (https://github.com/KCL-BMEIS/niftyreg). The minor modifications to SPM12 unified segmentation required to allow resumption of parameter estimation are available with the MCALT template. The Frangi filter implementation is available at https://www.mathworks.com/matlabcentral/fileexchange/24409-hessian-based-Frangi-vesselness-filter.

## Results

Results presented here are intended to evaluate the practicality and reasonableness of results across an age-stratified cohort. The cohort selection is not representative for clinical/biological interpretation beyond rudimentary comparisons.

Visual QC was performed for the 773 image sets. In one image set the segmentation model failed to converge to a plausible solution; this was detected by automated contrast checking and confirmed by visual assessment. A total of 16 image sets failed segmentation QC. Underlying reasons included extremely poor T1- and/or T2-weighted image quality generally due to patient motion, the presence of large strokes rendering the cortical anatomy uninterpretable, and the previously mentioned segmentation failure. Additionally, 10 image sets were eliminated due to PVS under-identification. The remaining image sets (n=747 or 96.7%) were retained for analysis. Data presented in Figures 2-6 was selected as follows: one point per participant is shown when values are plotted versus age; 300 subjects were randomly chosen for longitudinal “spaghetti” plots to reduce over-crowding.

**Figure 2.**
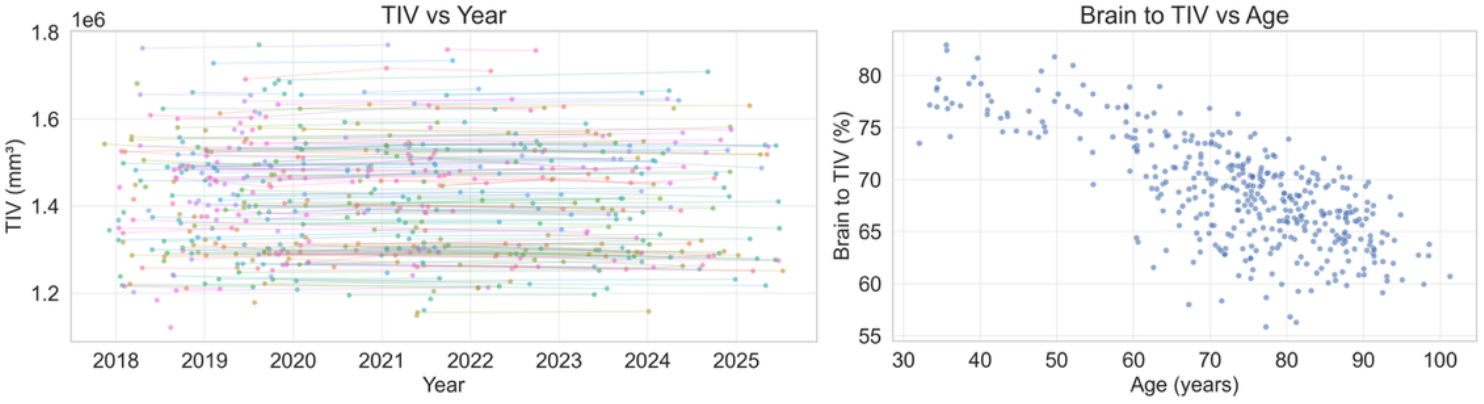
Total intracranial volume (TIV) versus scan date and brain volume are a percentage of TIV versus age. TIV versus image acquisition date in individuals with serial exams is shown in the left plot including longitudinal data. As the population contains adults, TIV is expected to be stable over time. In the right plot brain volume relative to TIV decreases with age.

**Figure 2.**
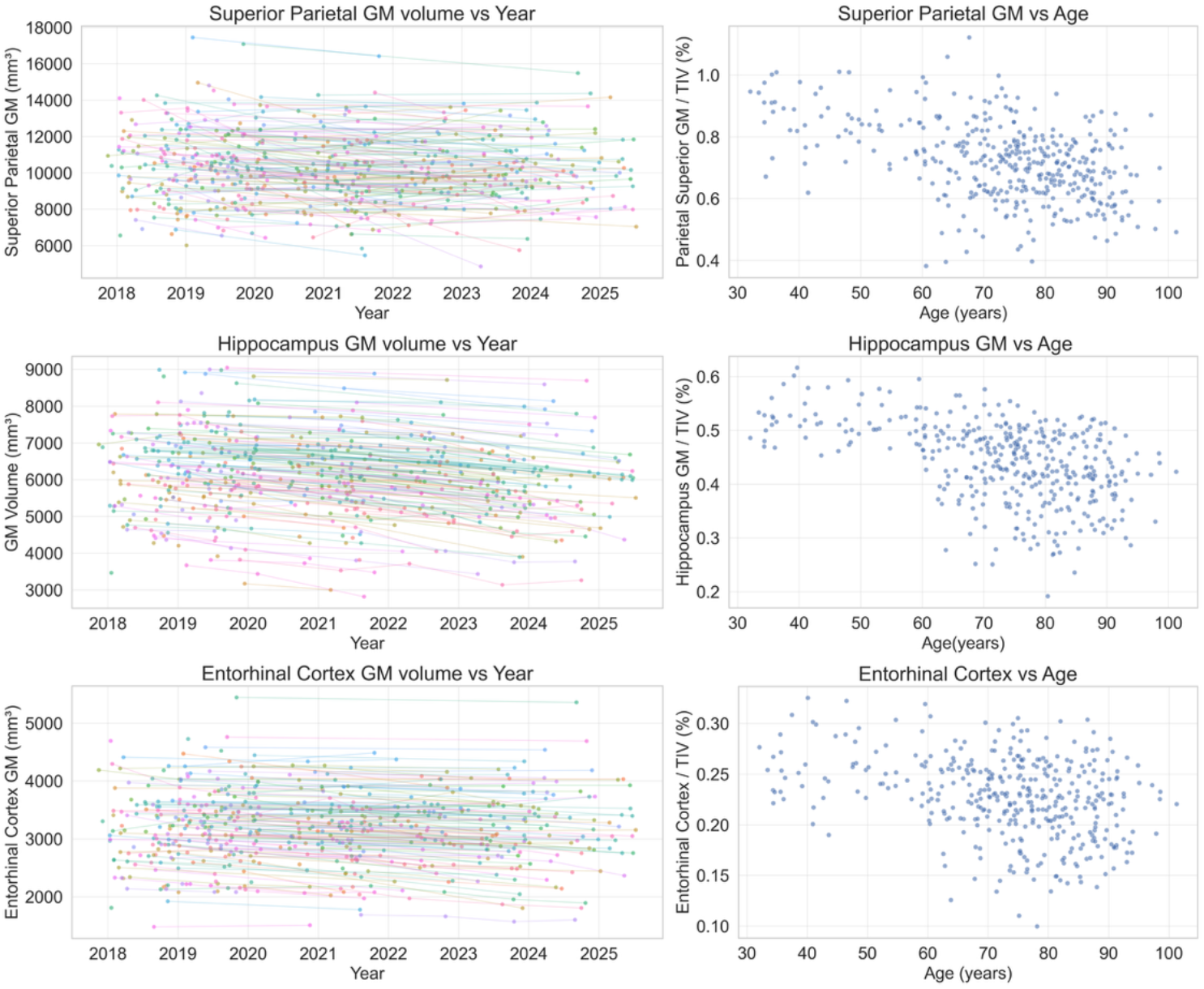
Superior Parietal, Hippocampal and Entorhinal Cortex GM volumes. GM volumes are stable to decreasing over time in individuals with serial exams. GM volume, normalized to TIV, tends to decrease with age.

TIV is assumed to be constant over time in adults. Multiple MRIs were analyzed for 249 participants. For each participant with multiple MRIs the deviations around the subject mean were calculated. The resulting distribution had a standard deviation of approximately 0.38%. For comparison, the variability introduced in scanner gradient calibration by routine service on Siemens scanners appears in discrete steps of one part in 512, or approximately 0.2% (Gunter et al., 2009). Each gradient axis is independently calibrated, and volume uncertainties include the product of the (unknown) calibration errors. Observed TIV variability is of the same order of magnitude as expected calibration-related uncertainty. The total brain tissue volume (TTV) was also calculated from the sum of voxels identified as GM, NAWM, DGM, and WMH. In Figure 2, TIV vs date (including longitudinal series) and the ratio of brain volume to TIV vs age are (one value per subject) shown. The range of brain volumes is approximately 1.2 to 1.8L. The brain to TIV ratio tends to decrease with age as expected.

Gray matter volumes are presented in Figure 3 and Figure 4. GM volumes are assessed using the 122-region atlas from MCALT. We selected two meta-regions for Figure 3, the supratentorial GM volume and an “AD Signature” region (Jack et al, 2015) that focuses on regions typically involved in AD, specifically the entorhinal cortices, fusiform gyri, inferior temporal gyri and the middle temporal gyri. In Figure 4, analogous plots are shown for GM in the superior parietal lobe, hippocampus and entorhinal cortex. The superior parietal region was chosen as a representative of the superior cortex. The hippocampus is a relatively large GM structure, of interest in AD. The entorhinal cortex is smaller and was chosen as it can be a challenging segmentation target. Again, longitudinal data are plausible – typically stable or decreasing volumes with time. Cross sectional values tend to decrease and demonstrate increasing variability with age.

**Figure 3.**
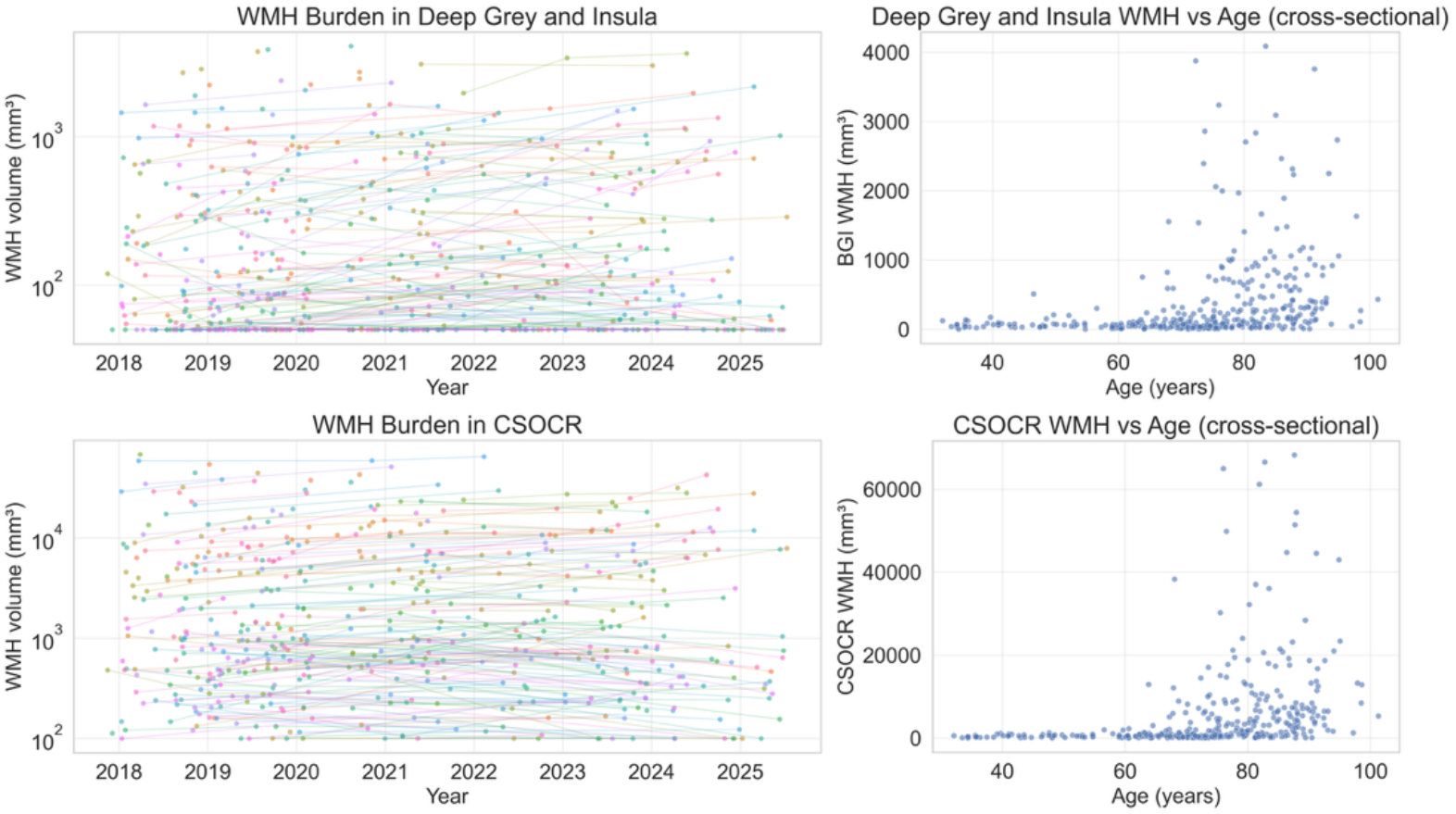
WMH burden in Deep Gray + Insula and CSOCR versus image acquisition date (left column) and versus age (right column). Longitudinal data is log() transformed to more clearly show trajectories of individuals. Furthermore, volumes below 50mm^3 and 100mm^3 are clipped for the Deep Gray and Insula and CSOCR regions respectively. Trajectories are generally increasing over time in individuals with serial exams.

**Figure 4.**
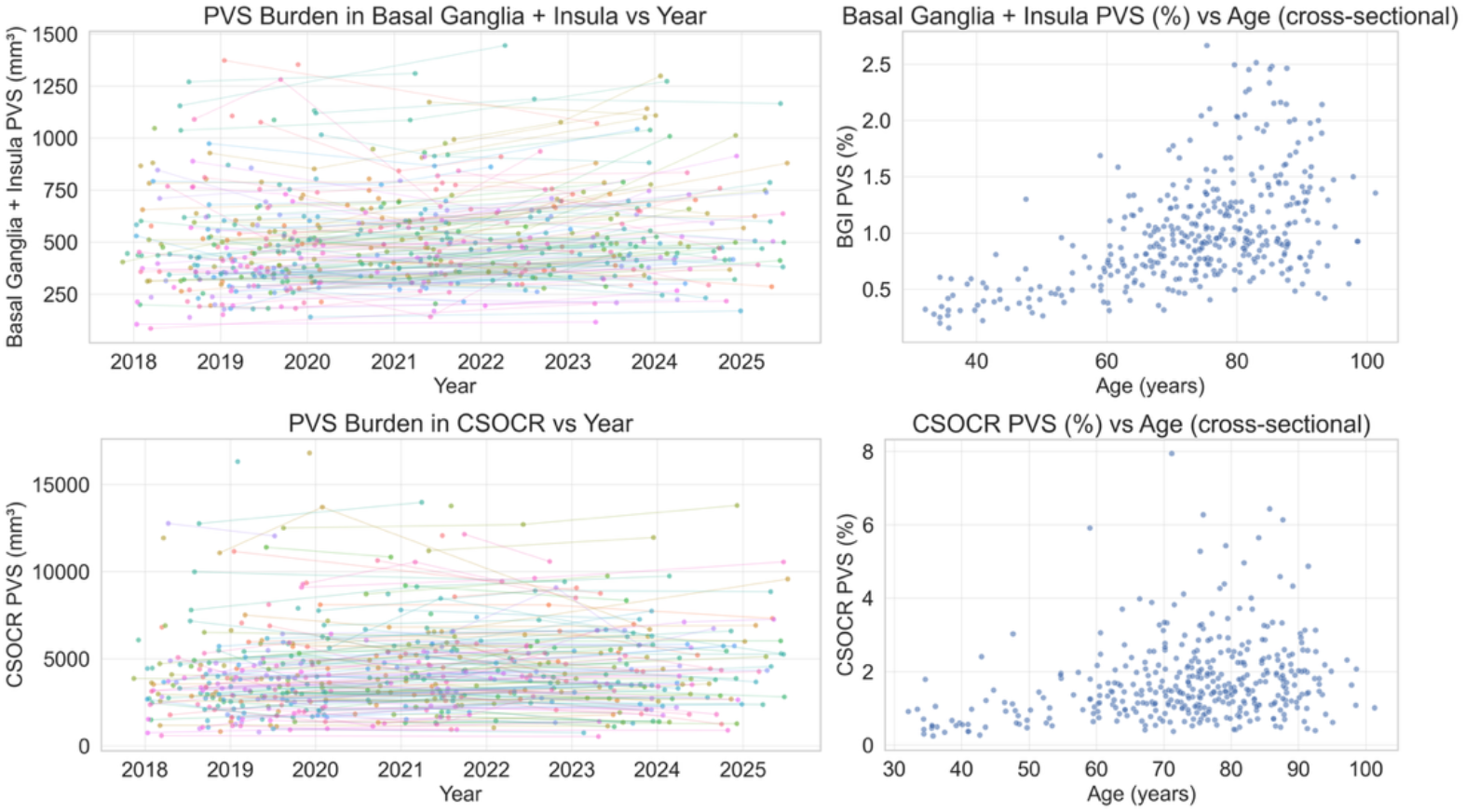
PVS burden in Deep Gray + Insula and CSOCR versus image acquisition date (left column) and versus age (right column). PVS burden is expected to generally increase with age. Longitudinal trajectories of PVS burden in basal ganglia and insula are flat to increasing over time in individuals with serial exams, while PVS burden trajectories appear more flat in the CSOCR

WMH and PVS results were derived using the 42-region lobar atlas from MCALT. For both WMH and PVS only voxels overlapping the PVS search space are considered here. An approximate centrum semiovale + corona radiata region (CSOCR) region is created by summing frontal, parietal, and occipital lobar atlas regions. The deep gray nuclei and insula are summed to create a “Basal Ganglia + Insula” region. WMH volume vs date and WMH volume vs age are shown in Figure 3. Figure 4 provides the same comparisons for PVS volume burden. Both WMH and PVS burden are expected to be higher in older participants and to exhibit stable longitudinal change over a period of years (DeCarli et al., 2005; Schmidt et al., 2003; Wardlaw et al., 2013; Ding et al., 2017).

The number of automated PVS clusters found is lower than by visual assessment. Visual assessment relies on single image slices, whereas automated PVS clustering is in 3D. Therefore, multiple visually counted PVS may be part of a larger cluster. One image set was removed from analysis in the basal ganglia where visual assessment found roughly 40 PVS and automated clustering approximately 20. Visual inspection of that image found relatively large areas which clustering combined and were counted as multiple PVS on visual assessment. Deming regression was performed to calibrate the automated result to the mean of visual counts across raters. Scatter plots after calibration and Bland-Altman plots are shown in Figure 5. Supplemental data Figure S1 includes all data with no calibration.

**Figure 5.**
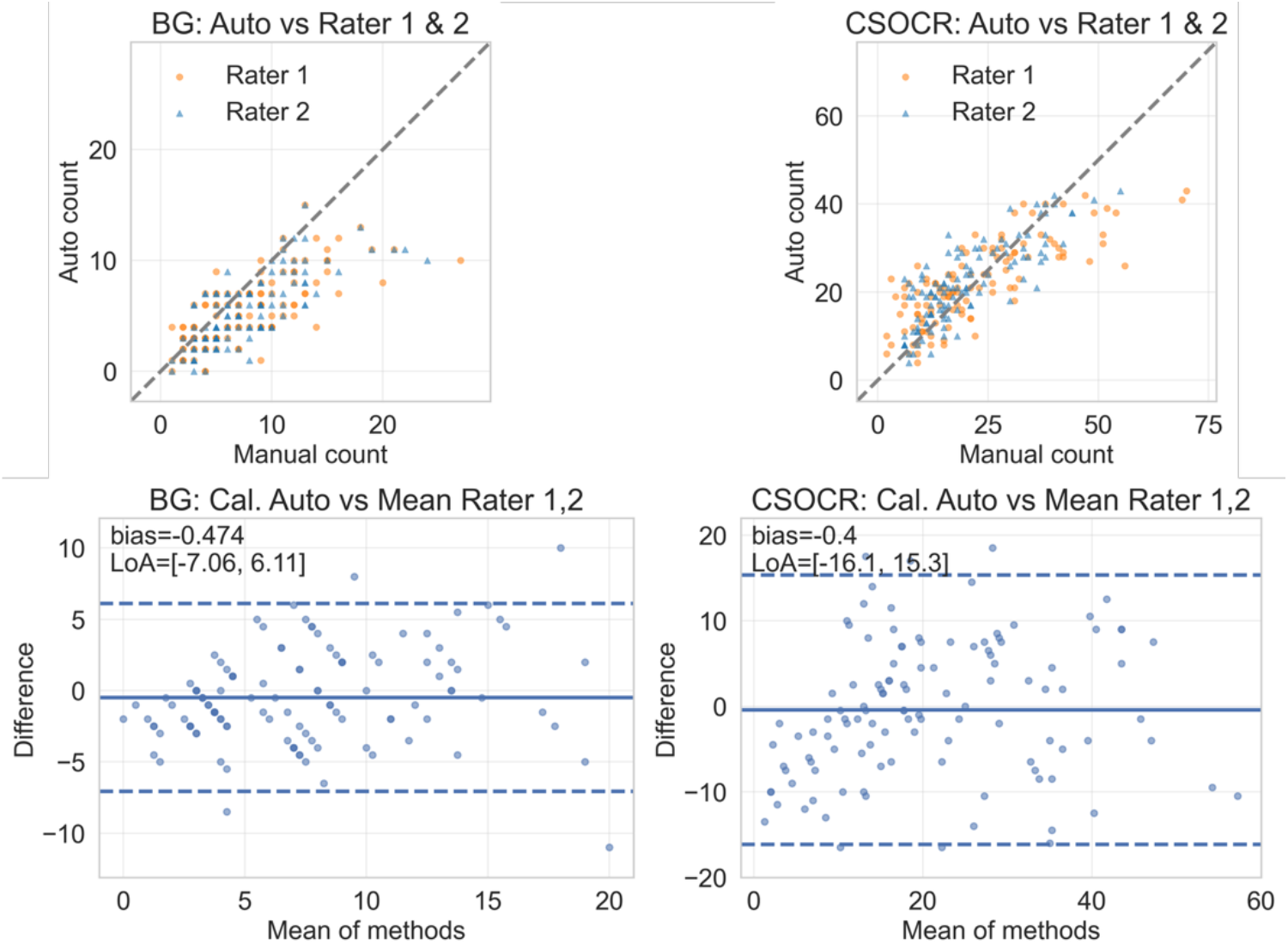
Comparison of calibrated auto counts to the mean of visual PVS assessment in the basal ganglia (BG) slice and centrum semiovale / corona radiata (CSOCR) slice. Scatterplots are presented in the top row with the identity line for reference; Bland-Altman plots with the 95% limits of agreement are shown in the bottom row. Calibration reduced the bias to near zero as expected.

## Discussion

Utilizing multiple MRI contrasts allowed us to extend probabilistic segmentation beyond gray-white-and-CSF, adding tissue classes for WMH and DGM. Using Frangi filtering on T1- and T2-weighted images, we created a per-subject prior of vessel-like shapes. This prior was substituted into the CSF channel of the previously estimated segmentation model to find PVS voxels. The resulting method was robust as observed by the following results: 1) Cross-sectional results as a function of age were consistent with expectations with TIV being stable across age, with brain tissue and GM decreasing with age, and the burden of cerebrovascular disease (WMH and PVS) increasing with age. 2) The longitudinal results were stable over time with trends as expected with age. While this work was focused on developing and evaluating the robustness of the method on a large age range, future work will focus on specific biological questions.

Tissue segmentation is the basis of MRI-based quantification of biomarkers. With most aging and dementia studies now acquiring multiple contrast images to assess brain health, the proposed methodology can be employed to study multiple aspects of brain health. While methods do exist for segmentation of WMH using T2-FLAIR MRI or in combination of T1 images, the simultaneous consideration of tissue classes enhances the robustness of our proposed methodology. This is reflected in the longitudinal analysis where the trends were stable and without outliers that go up and down even though each time point was processed independently.

PVS are thin and tubular structures that fall near or below the resolution of the MRI acquisition. The partial volume and low contrast make PVS segmentation a difficult problem. Automated methods have been proposed using classical image processing methods and deep learning methods for segmentation of PVS. The multi contrast approach we undertook here in combination with classical image processing methods provides a robust way to segment PVS. Two key findings support our methodology – strong agreement with manual ratings and stable longitudinal trends that are critical for studying mechanisms.

There are some limitations of this work. It requires multiple good quality image sets. Within this data, however, total losses (N=16 out of 773 or approximately 2%) were modest. The three image sequences used here require approximately 17 minutes of MRI scanning time, which is significant compared to a common 30- or 45-minute time slot for scanning. Each sequence is 5 to 8 minutes long. Typical aging and dementia studies acquire T1 and T2-FLAIR sequences for basic assessment of the brain. While acquisition of conventional T2-weighted images adds time and may be removed due to pressure to shorten protocols, the information available from the three channels is invaluable for robust PVS segmentation. We note, however, that the segmentation model itself can be applied using only T1-weighted and T2-FLAIR images for datasets that do not have T2-weighted images or for historical datasets provided that PVS identification is surrendered. While the use of imaging from a single scanner manufacturer with a range of scanner software versions is a limitation, we have found that protocols that are reasonably consistent in large aging and dementia studies such as ADNI (that acquire the three channels) will allow the utilization of this method.

Obvious future directions include assessing the PVS and WMH in models of cerebrovascular risk, aging, and various dementias. Comparisons with various single output models is warranted. Recent large-scale longitudinal work demonstrates that automated MRI-derived PVS metrics are associated with increased dementia risk and accelerated brain atrophy (Barisano et al., 2025). Finally, this provides a way to generate very large self-consistent training sets for AI model development.

## Data availability

Numeric data are available upon request. The expanded MCALT tissue priors will be included in the next MCALT update (https://www.nitrc.org/projects/mcalt/). Images from MCSA are available from the MCSA project on GAAIN.org, and those from the Mayo ADRC are available as part of the SCAN project https://scan.naccdata.org.

## Ethics Statement

All imaging data used in this study were acquired as part of research protocols approved by the Mayo Clinic Institutional Review Board (IRB). Written informed consent was obtained from all participants prior to participation, in accordance with the Declaration of Helsinki and applicable institutional and federal guidelines. The use of these data for the present analyses was covered under the original IRB approvals.

## Acknowledgments

The imaging and development was funded through US NIH grantsU01 AG006786 (PIs: Petersen, Jack, Graff-Radford, Vemuri), R01 AG056366 (PI: Vemuri), R37 AG011378 (PI: Jack), P50 AG016574 (PI: Petersen), R01 AG034676 (PI: Rocca); the GHR Foundation grant, the Alexander Family Alzheimer’s Disease Research Professorship of the Mayo Foundation, the Elsie and Marvin Dekelboum Family Foundation, U.S.A. and Opus building NIH grant C06 RR018898.

## Supplementary Materials

### Visual QC Approach

As described in the main text images are automatically loaded for QC based on filenames and contrast settings derived from the segmentation output file. The T1-weighted image window leveling is adjusted so that the mean NAWM intensity from the segmentation model is 60% of the maximum displayed intensity. The T2-weighted image window leveling is adjusted so that the mean NAWM intensity from the segmentation model is 45% of the maximum displayed intensity. The T2-FLAIR image window leveling is adjusted so that the mean WMH intensity from the segmentation model is 50% of the maximum displayed intensity. The PVS labels are displayed as an opaque light blue overlay. The argmax segmentation is color mapped and displayed at 40% opacity. TIV masks and atlas regions are colormapped and displayed as opaque overlays.

Multiple review passes are required as multiple outputs are generated. The initial pass procedes superior to inferior viewing the T2-FLAIR image until a WMH lesion is found, periventricular WMH is observed or, in rare cases, the bottom of the cerebellum is reached. This initial pass allows the evaluator to develop an overall sense for WMH load and patient anatomy. The semi-opaque argmax segmentation is enabled and the superior to inferior review is restarted at the top of the brain. In this step the GM, NAWM, and WMH classification is evaluated. Empirically, GM segmentation fails only when the patient neuroanatomy cannot be reasonably mapped to the template anatomy (e.g. the presence of a large stroke). Next, the T2-weighted image is inserted in place of the FLAIR image and a sagittal slice about 10mm off the mid-line is checked to confirm that CSF segmentation extends to the dura and that the CSF/GM/NAWM borders appear correct. The argmax segmentation is turned off and the evaluator checks for PVS in the CSOCR to calibrate expectations for PVS load. The PVS overlay is then enabled. PVS assessment is carried out evaluating sagittal slices, nominally from right to left. The PVS overlay may be toggled if the evaluator questions mask correctness. Furthermore, the T1-weighted image may be consulted. In cases with high BG PVS load a second left to right pass may be necessary. Finally the atlas and TIV overlays are checked with the T1-weighted image as a background.

### Visual Assessment Implementation

The Potter method for visual assessment includes selection of a single slice of a T2-weighted image in the basal ganglia and a single slice in the centrum semiovale / corona radiata and counting the number of bright clusters in the hemisphere with the most clusters. Knowing that we would be comparing automated counts and visual counts we sought to optimize consistency in the visual assessments. For each MR study to be assessed, the T1-weighted image was aligned (6DOF) to the MCALT gray scale average image and single channel gray-white-CSF segmentation performed. The T2-weighted image was registered to the aligned T1-weighted image and resampled into that voxel raster. Voxels identified as predominantly white matter were identified from the segmentation output. In those voxels, T2 voxel intensities were identified. The contrast was adjusted so that the dynamic range spanned from zero to 2.5 times the mean value. Each rater could adjust the contrast, but this provided a consistent starting point. Slice and hemisphere selection was done by consensus review of the raters. Raters marked each PVS by placing a dot in the appropriate location. Dots were automatically counted. Automated clusters in the same regions were counted. Empirically, human raters often separated bright regions that would be clustered together into multiple dots based on local intensity maxima.

### MRI Acquisitions

Structural MRI acquisitions spanned two protocol epochs. All images were acquired on a Siemens Prisma Scanner using a 64-channel head coil. The earlier epoch, through early 2020, included a 3D T1-weighted MPRAGE (0.8 mm isotropic; TR/TE/TI ≈ 2300/3.14/945 ms, flip angle 9°), a 3D T2-weighted sequence (0.8 mm isotropic; 3D T2-SPACE, TR/TE ≈ 3200/409 ms), and a sagittal 3D T2-FLAIR with slightly anisotropic resolution (≈1.0 × 1.0 × 1.2 mm; TR/TE/TI ≈ 4800/441/1650 ms). The later epoch followed the ADNI-4 structural MRI protocol which specifies 1.0 mm isotropic 3D acquisitions for all core structural contrasts. For Siemens Prisma implementations, ADNI-4 specifies a 3D T1-weighted MPRAGE (TR/TE/TI ≈ 2300/2.98/900 ms, flip angle 9°), a sagittal 3D T2-FLAIR (TR/TE/TI ≈ 4800/441/1550 ms), and a 3D T2-weighted sequence (SPACE; TR/TE ≈ 3200/409 ms).

**Figure S.**
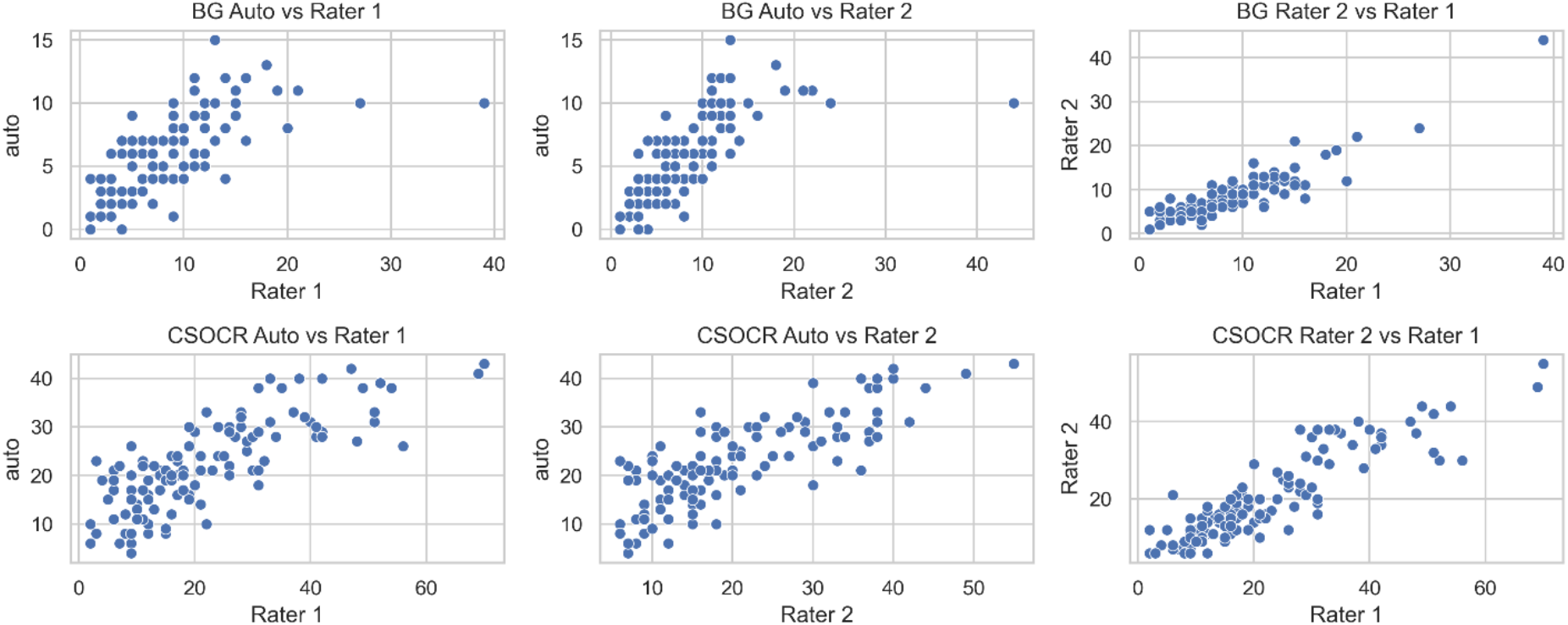
Comparison of automated and visual PVS counts. “auto” refers to the number of PVS clusters found in the same slice and hemisphere as was visually assessed. Counts from the Basal Ganglia slice are shown in the top row. Counts from the CSOCR are shown in the bottom row. The outlier with approximately 40 visually counted PVS is shown here but excluded in the Results section.

